# Transcriptome analysis of the Molecular Mechanism underlying Immunity- and Reproduction trade-off in *Locusta migratoria* Infected by *Micrococcus luteus*

**DOI:** 10.1101/524611

**Authors:** Shaohua Wang, Xiaojun Liu, Zhiyong Xia, Guoqiang Xie, Bin Tang, Shigui Wang

**Affiliations:** Hangzhou Key Laboratory of Animal Adaptation and Evolution, College of Life and Environmental Sciences, Hangzhou Normal University, Hangzhou, Zhejiang, 310036, China

**Keywords:** transcriptome, immune defense, reproduction, trade-off, *Locusta migratoria*, *Microcroccus luteus*

## Abstract

Immune response and reproductive success are two of the main energy-consuming processes in living organisms. However, it is unclear which process is prioritized when both are required. Therefore, the present study was designed to examine this question using one of the world’s most destructive agricultural pests, the migratory locust *Locusta migratoria.* Transcripts from the ovaries and fat bodies of newly emerged locusts were analyzed, using RNA-seq based transcriptome and qualitative real-time PCR, at 4 h and 6 d after being infected with the gram-positive bacteria *Microcroccus luteus,* and changes in the main biological pathways involved in reproduction and immunization were analyzed using bioinformatics. At 4 h after infection, 348 and 133 transcripts were up- and down-regulated, respectively, whereas 5699 and 44 transcripts were up- and down-regulated, respectively, at 6 d after infection. Meanwhile, KEGG analysis indicated that vital pathways related with immunity and reproduction, such as Insulin resistance, FoxO signaling, Lysosome, mTOR signaling, and Toll-like receptor signaling pathways were up-regulated. Among the differentially expressed genes, 22 and 17 were related to immunity and reproduction, respectively, and the expression levels of *PPO1* and *antimicrobial peptide defensin 3* were increased (log_2_FC = 5.93 and 6.75, respectively), whereas those of *VgA* and *VgB* were reduced (log_2_FC = −17.82 and −18.13, respectively). These results indicated that that locusts allocate energy and resources to maintain their own survival by increasing immune response when dealing with both immune and reproductive processes. The present study provides the first report of expression levels for genes related with reproduction and immunity in locusts, thereby providing a reference for future studies, as well as theoretical guidance for investigations of locust control.

## Introduction

Insects are the most successful group of animals on earth, owing, at least in part, to the effectiveness of their immune response to microbial invasion, and the immune responses of insects to pathogenic infection include both humoral and cellular immunity. Humoral immunity mainly includes the synthesis of antimicrobial peptides (AMPs) in the fat body and is primarily based on the Toll and Imd signaling pathways, which play important roles in AMP expression, as well as on the interaction between the two pathways (Ferrandon et al., 2007; Viljakainen L, 2015). Meanwhile, cellular immunity includes hemocyte-mediated phagocytosis, encapsulation, and nodulation. The melanization of macroparasites triggered by hemocytes participating in phagocytosis or phenoloxidase cascade activation are hallmarks of cellular response (Kirschman LJ et al., 2017). Furthermore, the JNK and JAK/STAT pathways also contribute to the immune response (Uvell H et al., 2007). During these immune processes, insects must invest energy and resources to survive against a variety of pathogens.

Recently, the reproduction process of female insects has been thoroughly studied as an important target for pest control (Roy S et al., 2017). The process is mainly regulated by juvenile hormones (JH), ecdysteroids, and nutritional signaling pathways, and differs among insect species owing to differences in reproductive strategies (Raikhel AS et al., 2005). In most insect species, the central part of female reproduction is vitellogenesis, which includes the production of both vitellogenin (Vg) and other yolk protein precursors (YPPs), followed by the internalization of YPPs by maturing oocytes *via* receptor-mediated endocytosis (Swevers L et al., 2005). Depending on tissue type, sex, and developmental stage, Vg is synthesized extra-ovarially by the fat body, secreted into the hemolymph, and then sequestered by competent oocytes *via* receptor-mediated endocytosis (Swevers L et al., 2005). Vitellogenin is stored in a crystalline form, as vitelline, after being incorporated into oocytes, where it functions as a reserve food source for future embryos (Tufail M et al., 2008). The reproductive process may be affected by energy metabolism, nutrient metabolism, and fitness value. In *Locusta migratoria* (Orthoptera), female reproduction is governed by JHs, which contribute to vitellogenesis and oocyte maturation (Luo M et al., 2017).

Although both reproduction and the ability to survive pathogenesis are essential functions, the evolution of life history is a matter of optimization, rather than maximization, so that organisms must partition limited energy and nutritional resources among a variety of life processes. The trade-off between female insect reproduction and immunity has been reported in recent years (Schwenke RA et al., 2016). Because both immunity and reproduction are physiological energy-consuming processes, increasing reproduction effort reduces immune ability, whereas immune system activation reduces reproductive output (Stearns SC, 2000). The locust is one of the most important agricultural pests worldwide. Based on the study of the trade-off between immunity and reproduction, it is interesting to identify genes that are involved in both reproduction and immunity, to clarify the internal regulatory mechanism of such genes, and to identify the best biological target for pest control. However, little research has been conducted on the trade-off between immunity and reproduction in locusts. In the present study, the gram-positive bacteria *Micrococcus luteus* was used to infect newly emerged locusts. The fat body and ovary, which are the main organs involved in immunity and reproduction, were dissected at 4 h and 6 d after parasitism.

Differentially expressed genes (DEGs) analysis based on next-generation RNA-seq technology provides extensive data, with enormous depth and coverage, thereby allowing the global analysis of a parasitized host’s transcription profile. To identify immunity- and reproduction-related genes that were differentially up- or down-regulated in the infected locusts, the present study compared the fat body and ovary transcripts of (1) the control and infected locusts at 4 h after treatment, (2) the control and infected locusts at 6 d after treatment, (3) the control locusts at 4 h and 6 d after treatment, and (4) the infected locusts at 4 h and 6 d after treatment. Bioinformatic analysis and qRT-PCR verification were used to predict and explore genes involved in immunity and reproduction, as well as to identify the molecular mechanisms underlying energy and resources trade-off in *L. migratoria.*

## Materials and Methods

### Insect Rearing and Experimental Treatments

Eggs were collected from *L. migratoria* that were maintained in the insect laboratory of Hang Zhou Normal University (Zhejiang Province, China). The locust models were established by rearing the insects in each well-ventilated cage (50 × 50 × 50 cm) at densities of 200–300 insects per cage. The insects were reared using a 16 h photoperiod at 30 ± 1°C and were fed a diet of fresh greenhouse-grown wheat seedlings (Ma Z et al., 2011; Wu R et al., 2012). Female locusts were collected and subject to parasite treatments within 12 hours after eclosion. The treated group was infected with the bacterial pathogen *M. luteus* (American Type Culture Collection, Manassas, VA, USA), which had been cultured to the logarithmic phase, according to the manufacturer’s instructions, by injecting the pronotum of each locust with 10 μl of bacterial culture (0.9 optical density at 600 nm) using a nanofil syringe (Shenggong, Shanghai, China). Meanwhile, the pronotum of each locust in the control group was stabbed using an alcohol-sterilized needle. For both the treatment and control groups, the wounding site of each insect was sealed using petroleum jelly to prevent exogenous natural infection. A diet of wheat seedlings and wheat bran was supplied in sufficient, but not excessive, amounts. Fat bodies and ovaries were collected from females in each group at 4 h and 6 d after injection, respectively, with two biological replicates per group. Furthermore, 30 ovaries from females in each group were dissected and weighed.

### cDNA Library Construction and High-Throughput Transcriptome Sequencing

Trizol (Invitrogen, Los Angeles, CA, USA) was used to extract total mRNA from mixed tissue samples (fat body and ovary) taken from individuals from the control and treated groups and each time point us ing, following the manufacturer’s procedure. The quantity and purity of the total RNA were measured using a Bioanalyzer 2100 and RNA 6000 Nano LabChip Kit (RIN>7.0; Agilent, CA, USA). Poly(A) mRNA was isolated from ~10 μg of each total RNA sample using poly-T oligos attached to magnetic beads (Thermo-fisher, MA, USA). The isolated mRNA was fragmented using divalent cations under elevated temperature and then used to construct a cDNA library, following the protocol for the Illumina RNA ligation-based method (Illumina, San Diego, CA, USA). Briefly, the fragmented RNA was dephosphorylated at the 3’ end using phosphatase and phosphorylated at the 5’ end using polynucleotide kinase. These samples were purified using the RNeasy MinElute Kit (Qiagen, Dusseldorf, Germany), following the manufacturer’s instructions, ligated to a pre-adenylated 3’ adapter, which enabled the subsequent ligation of a 5’ adapter, and then subject to both reverse transcription and PCR. The average insert size for the paired-end libraries was 300 bp (±50 bp). Finally, the library was subject to single-end sequencing using an Illumina Hiseq 2000 by LC Sciences (Houston, TX, USA), following the vendor’s recommended protocol.

### Bioinformatic analysis

Raw data was filtered using Trimommatic software with default parameters. The clean reads that were mapped to the *L. migratoria* genome database (http://locustmine.org/index.html) were annotated using Locust Base-locust genome data, GO, KEGG, Nr, Nt, and Swissport. The number of perfect clean reads corresponding to each gene was calculated and normalized to the number of Reads Per Kilobase of exon model per Million mapped reads (RPKM) (Mortazavi A et al., 2008). Based on expression levels, DEGs were identified, using a P-value of ≤0.05 and log_2_ fold-change (log_2_FC) of ≥1. Gene ontology (GO) analysis (ftp://ftp.ncbi.nih.gov/gene/DATA/gene2go.gz) was conducted for functional classification of the DEGs, and pathway analysis was performed using KEGG (http://www.genome.jp/kegg).

### Quantitative Real-Time PCR Analysis

Fifteen genes were selected to verify the RNA-seq data. Primer 5.0 was used to identify appropriate primers (Sup Table 1), and the expression levels of the selected genes were normalized using the expression of actin. Quantitative real-time PCR (qRT-PCR) was performed using a Bio-Rad CFX96 Real-Time PCR Detection system (Bio-Rad, CA, USA) and Premix Ex Taq (SYBR Green) reagents (Takara, Dalian, China). The 20-μl qRT-PCR reactions were subject to a thermal profile of 95 °C for 3 min and then 40 cycles of 95 °C for 10 s and 60 °C for 30 s. The thermal melting profile was assessed using a final PCR cycle of 95 °C for 30 s, with temperature increasing constantly from 60 to 95 °C. Relative gene expression levels were calculated using the 2^-△△CT^ method, with three replicates per sample.

## Results

### Changes of ovaries in infected group and RNA-seq Library Analysis

We compared the weight of ovaries in infected group with control group. The weight of ovaries in infected group was significantly lighter than in control group (Fig 1). In RNA-seq experiment, we obtained single end reads with the quality high enough for the gene expression analysis subsequently (Table 1). In order to investigate patterns of gene expression related to locust immunity and reproduction, we analyzed control and infected groups at two time points, with two biological replicates each (eight samples in total). The *L. migratoria,* Nr, Nt, SwissProt, GO, and KEGG databases were used to annotate the transcripts (Table 2). Most of the mRNAs had low RPKM values, typically between 30 and 80 (Table 3).

**Fig 1.**
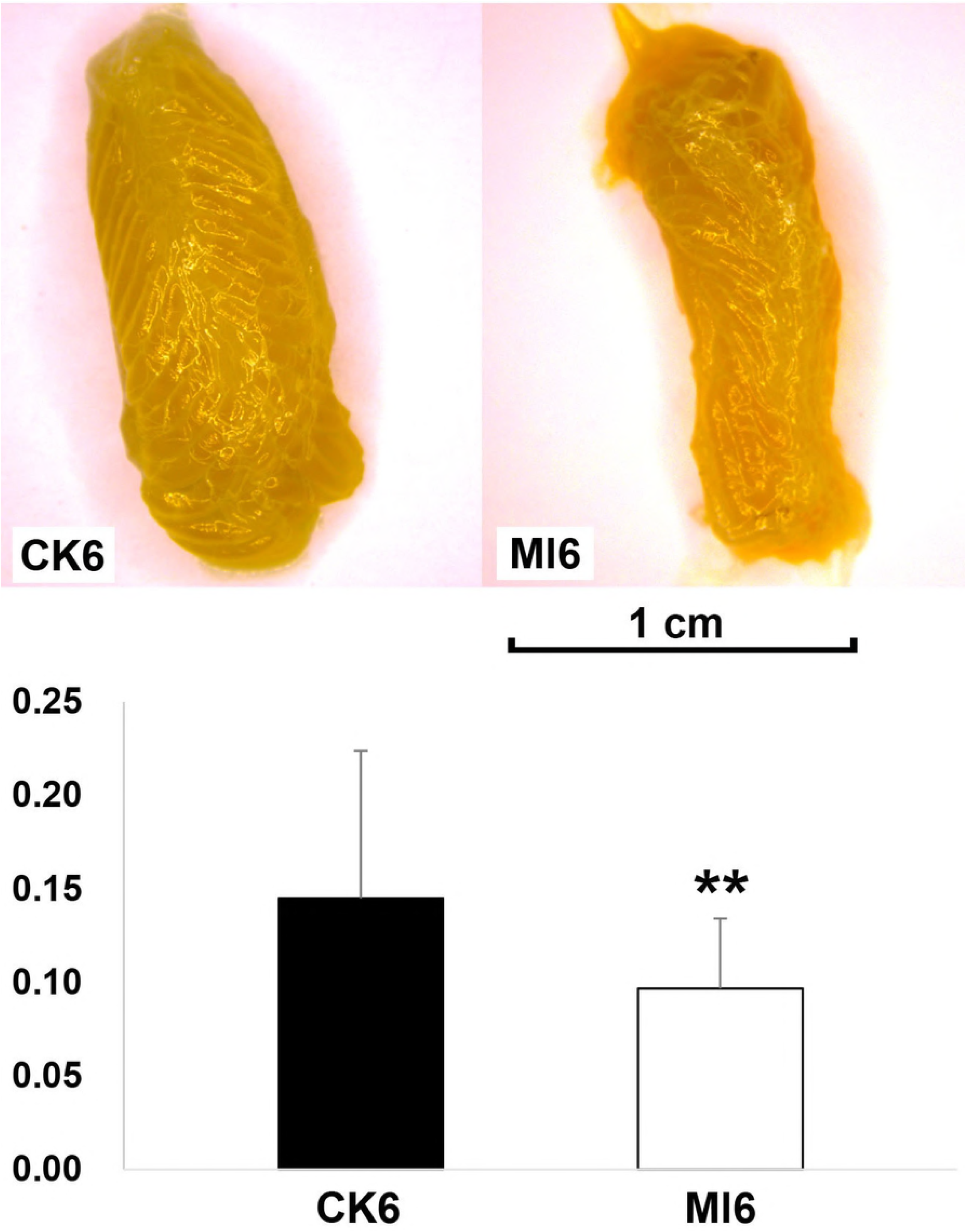
Comparison of ovary weight between control and infected group. Ml6 indicated the treatment group at 6 d after infection and CK6 indicated its control group.

**Table 1.** Overview of sequencing data.

| Sample name | Raw reads | Clean reads | Clean bases | Error(%) | Q20(%) | Q30(%) | GC(%) |
| --- | --- | --- | --- | --- | --- | --- | --- |
| CK4A | 14583856 | 14573380 | 0.73G | 0.02 | 99.37 | 97.67 | 43.19 |
| CK4B | 15370335 | 15359428 | 0.77G | 0.02 | 99.41 | 97.87 | 42.50 |
| CK6A | 18789172 | 18782102 | 0.94G | 0.03 | 99.37 | 97.69 | 49.17 |
| CK6B | 21062414 | 21058652 | 1.05G | 0.02 | 99.35 | 97.57 | 50.20 |
| Ml4A | 12436735 | 12228854 | 1.22G | 0.03 | 97.90 | 93.43 | 43.17 |
| Ml4B | 10058876 | 9885731 | 0.99G | 0.02 | 98.29 | 94.36 | 43.88 |
| Ml6A | 14433467 | 14199926 | 1.42G | 0.03 | 97.88 | 93.30 | 43.62 |
| Ml6B | 9681097 | 9492635 | 0.95G | 0.02 | 98.30 | 94.42 | 43.68 |

*Note: Sample: sample name; Raw Reads: four lines as a unit to statistic the sequence numbers of each document; Clean reads: filtered raw reads; Clean bases: number of sequences\* the length of sequence then turn to G; Error: mistake ration of base pairs; Q20、Q30: The percentage of bases whose Phred value is greater than 20,30 in all base pairs; GC content: The percentage of G and C in all base pairs.*

**Table 2.**
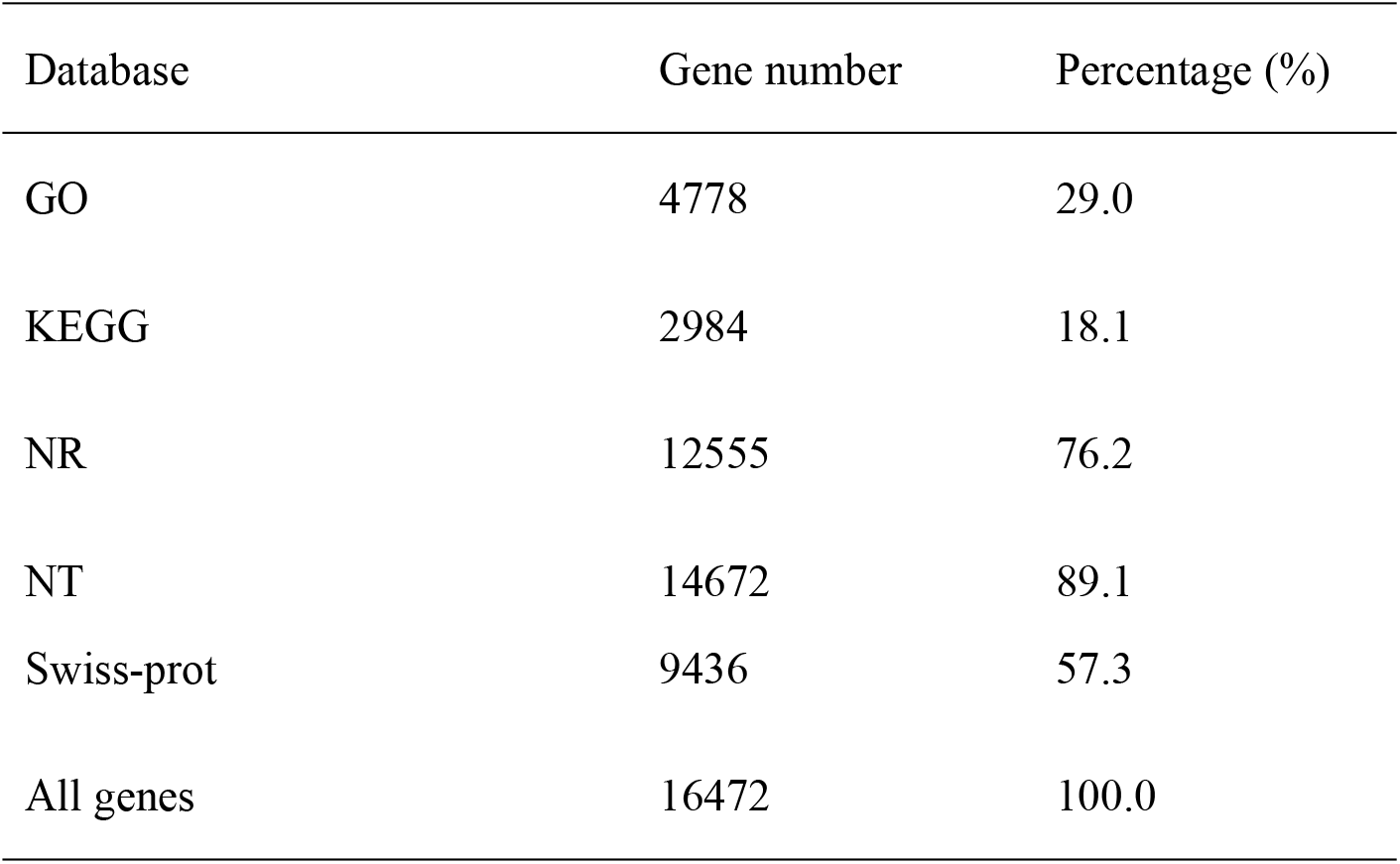
Annotation of identified genes.

| Database | Gene number | Percentage (%) |
| --- | --- | --- |
| GO | 4778 | 29.0 |
| KEGG | 2984 | 18.1 |
| NR | 12555 | 76.2 |
| NT | 14672 | 89.1 |
| Swiss-prot | 9436 | 57.3 |
| All genes | 16472 | 100.0 |

**Table 3.** RPKM expression analysis.

| Sample | Exp gene | Min. | Median | Mean | Max. | Sd. | Sum. |
| --- | --- | --- | --- | --- | --- | --- | --- |
| CK4A | 12888 | 0.01 | 11.98 | 80.75 | 104051.71 | 1035.33 | 1040761.9 |
| CK4B | 13862 | 0.01 | 21.79 | 64.36 | 15116.55 | 208.81 | 892134.56 |
| CK6A | 12670 | 0.01 | 3.25 | 31.37 | 66286.24 | 848.92 | 397499.26 |
| CK6B | 11877 | 0.01 | 1.56 | 30.69 | 72345.81 | 945.36 | 364532.7 |
| MI4A | 14121 | 0.01 | 20.69 | 64.31 | 3600.63 | 144.12 | 908109.58 |
| MI4B | 14003 | 0.01 | 20.77 | 64.68 | 2376.94 | 144.4 | 905717.15 |
| MI6A | 14411 | 0.01 | 19.05 | 66.32 | 17919.79 | 251.71 | 955735.42 |
| MI6B | 14247 | 0.01 | 20.04 | 67.31 | 7496.22 | 195.61 | 958994.88 |

*Note: Sample: sample name; Exp gene: genes identified in each sample; Min: minimum expression value of all genes in each sample; Median: median expression value of all genes in each sample; Mean: average expression value of all genes in each sample; Max: maximum expression value of all genes in each sample; Sd: variance of expression value of all genes in each sample; Sum: total expression value of all genes in each sample.*

### DEGs Analysis

First, the infected and control libraries for each time point were compared (Ml4 vs CK4 and Ml6 vs CK6) to identify genes that are differentially expressed at different times of infection. At 4 h after treatment, 348 and 133 genes were up-and down-regulated, respectively, in the infected group (Sup Table 2), and at 6 d after treatment, 5699 and 44 were up- and down-regulated (Sup Table 3). Secondly, the times points for the control and infected libraries were compared (CK6 vs CK4 and Ml6 vs Ml4) to identify genes that are differentially expressed during infection. For the control group, 24 and 1406 genes were up- and down-regulated, respectively, at 6 d after treatment, when compared to 4 h after treatment (Sup Table 4), and for the treatment group, 146 and 48 were up- and down-regulated (Sup Table 5). The number of the DEGs was summarized as columnar (Fig 2). All the DEGs were clustered to produce heat maps, the two replications of controls and treatments were clearly divided into two main clusters (Fig 3).

**Fig 2.**
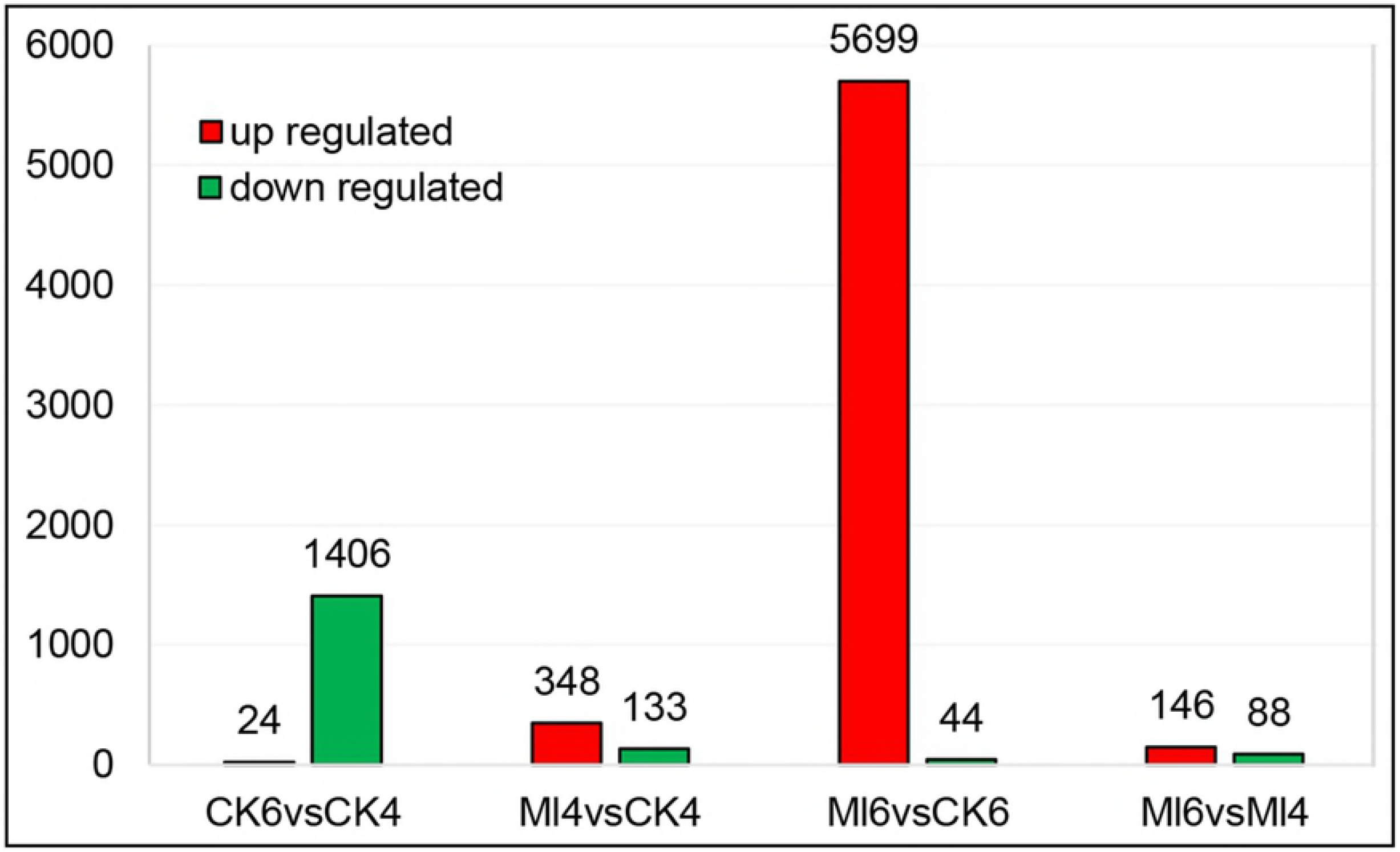
Transcripts that were significantly affected by either bacterial infection or time after treatment. Red and green indicate up- and down-regulated genes, respectively.

**Fig 3.**
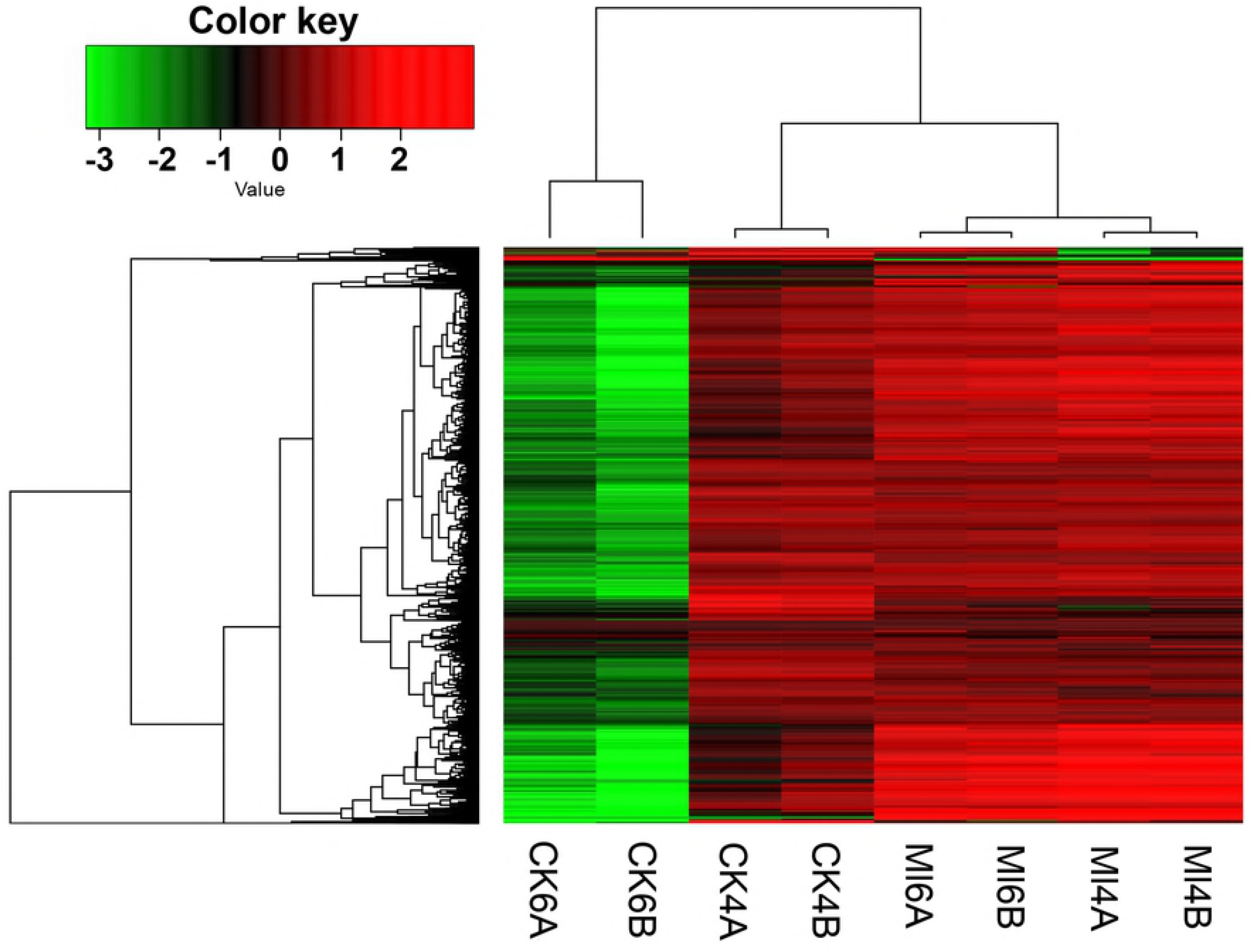
Clustering of differentially expressed genes.

### GO and KEGG Analysis

For the up-regulated GO terms related to immunity (Fig 4A, Sup Table 6), the most frequently mapped transcripts were ‘catalytic activity’ and ‘response to oxidative stress.’ In addition, a large number of transcripts were also mapped to functional groups related to fighting against foreign infections, such as ‘Lysosome’ and ‘Endosome’. ‘Immunity signal JNK cascade,’ ‘innate immune response,’ ‘defense response,’ ‘immune response,’ and ‘inflammatory response’ were represented by seven, seven, six, five, and three transcripts, respectively. For up-regulated GO terms related to reproduction, six GO terms were significantly changed, including ‘ovarian follicle cell development,’ ‘ovarian follicle cell stalk formation,’ ‘MCM complex,’ ‘ovarian nurse cell to oocyte transport,’ ‘female germline ring canal formation,’ and ‘negative regulation of TOR signaling.’ For GO analysis of the transcripts up-regulated at 6 d in the control group, most of the DEGs were related to genes involved in individual growth (Fig 5A). However, when comparing transcripts from the control and infected groups at 4 h after treatment and when comparing transcripts from the infected groups at 4 h and 6 d after treatment, none of the significantly changed GO terms were directly related to immunity or reproduction.

For the up-regulated transcripts mapped pathway (Fig 4B, Sup Table 7), 10 of them were involved in immunity and reproduction by the screening criteria of number of mapped transcripts was more than three: ‘Endocytosis,’ ‘Phagosome,’ ‘FoxO signaling pathway’ (Fig 6), ‘Lysosome’ (Sup Fig 1), ‘Insulin resistance’ (Fig 7), ‘mTOR signaling pathway’ (Sup Fig 2), ‘Regulation of autophagy,’ ‘Toll-like receptor signaling pathway,’ ‘Insulin signaling pathway,’ and ‘Insect hormone biosynthesis.’ At 4 h after treatment, the ‘Phagosome’ pathway (Sup Table 8) was up-regulated in the infected group. In the control group, five down-regulated pathways were identified as involved in immunity and reproduction, including ‘Endocytosis,’ ‘Lysosome,’ ‘Phagosome,’ ‘FoxO signaling pathway,’ ‘mTOR signaling pathway.’ (Fig 5B, Sup Table 9). In the infected group, no pathways related to immune or reproduction were differentially expressed at 6 d after treatments, when compared to 4 h after treatment (Sup Table 10).

**Fig 4.**
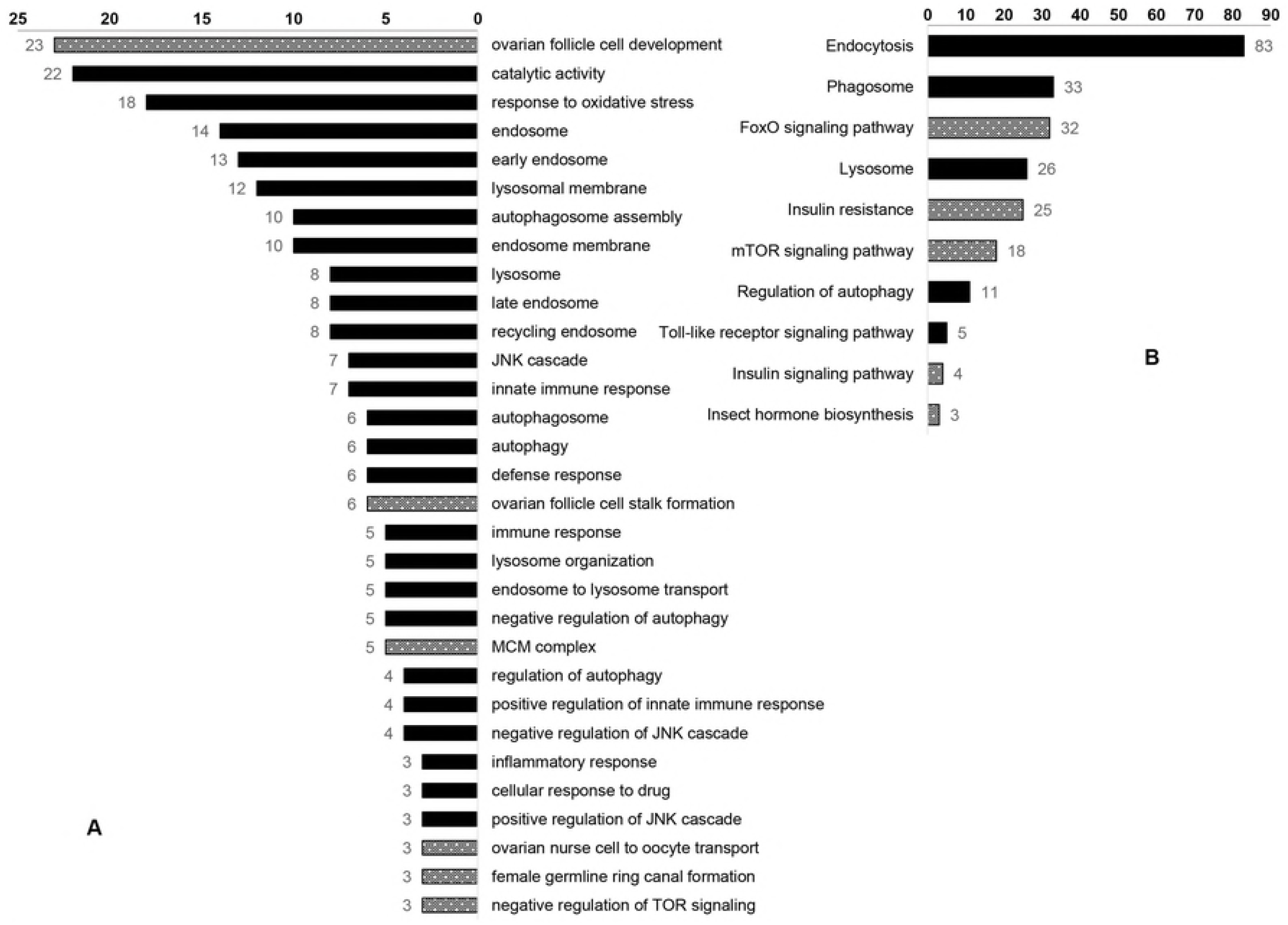
GO and KEGG terms for immunity- and reproduction-related genes that were differentially expressed among treatment groups at 6 d after infection.

**Fig 5.**
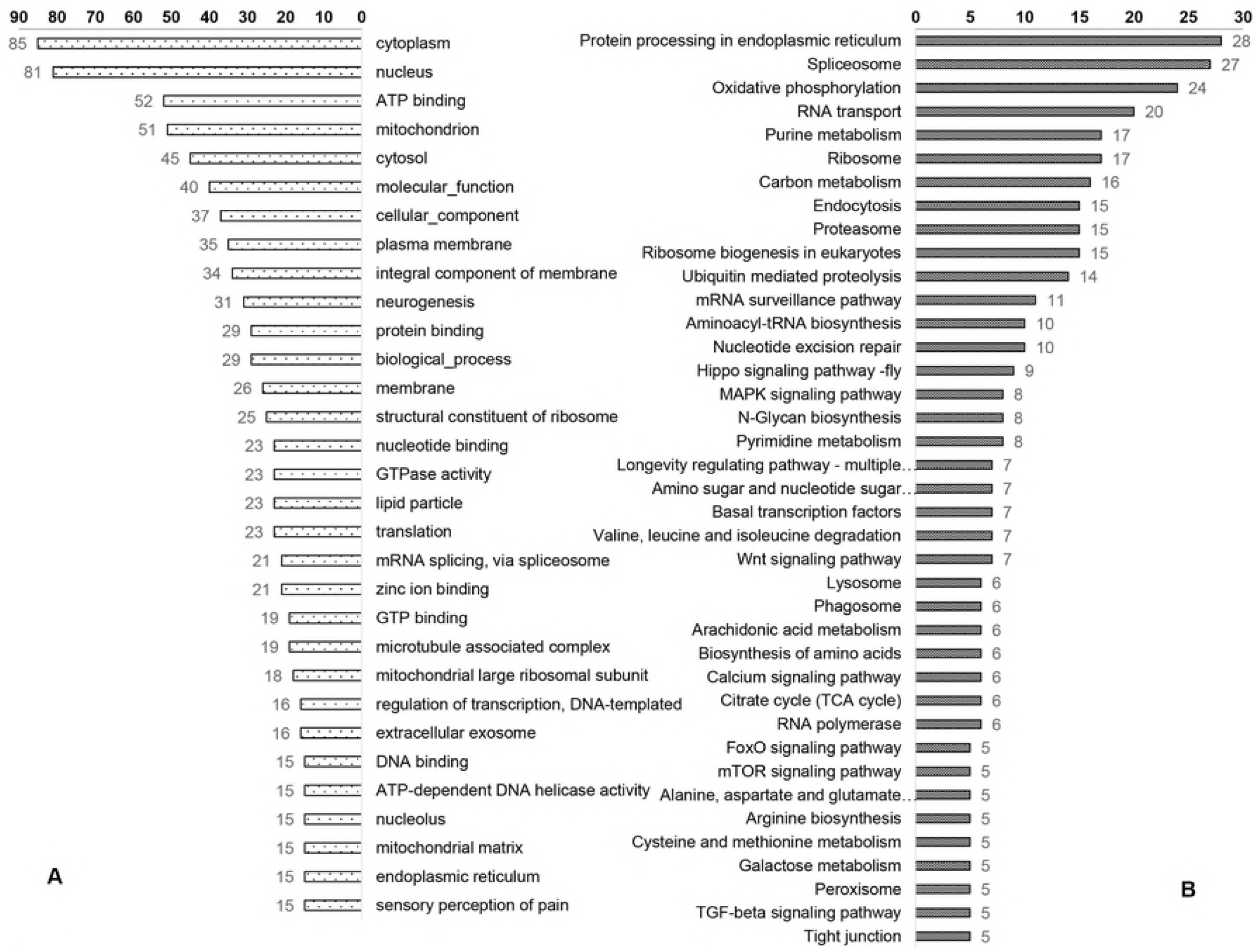
GO and KEGG terms for immunity- and reproduction-related genes that were differentially expressed in the control group at 4 h and 6 d after infection.

**Fig 6.**
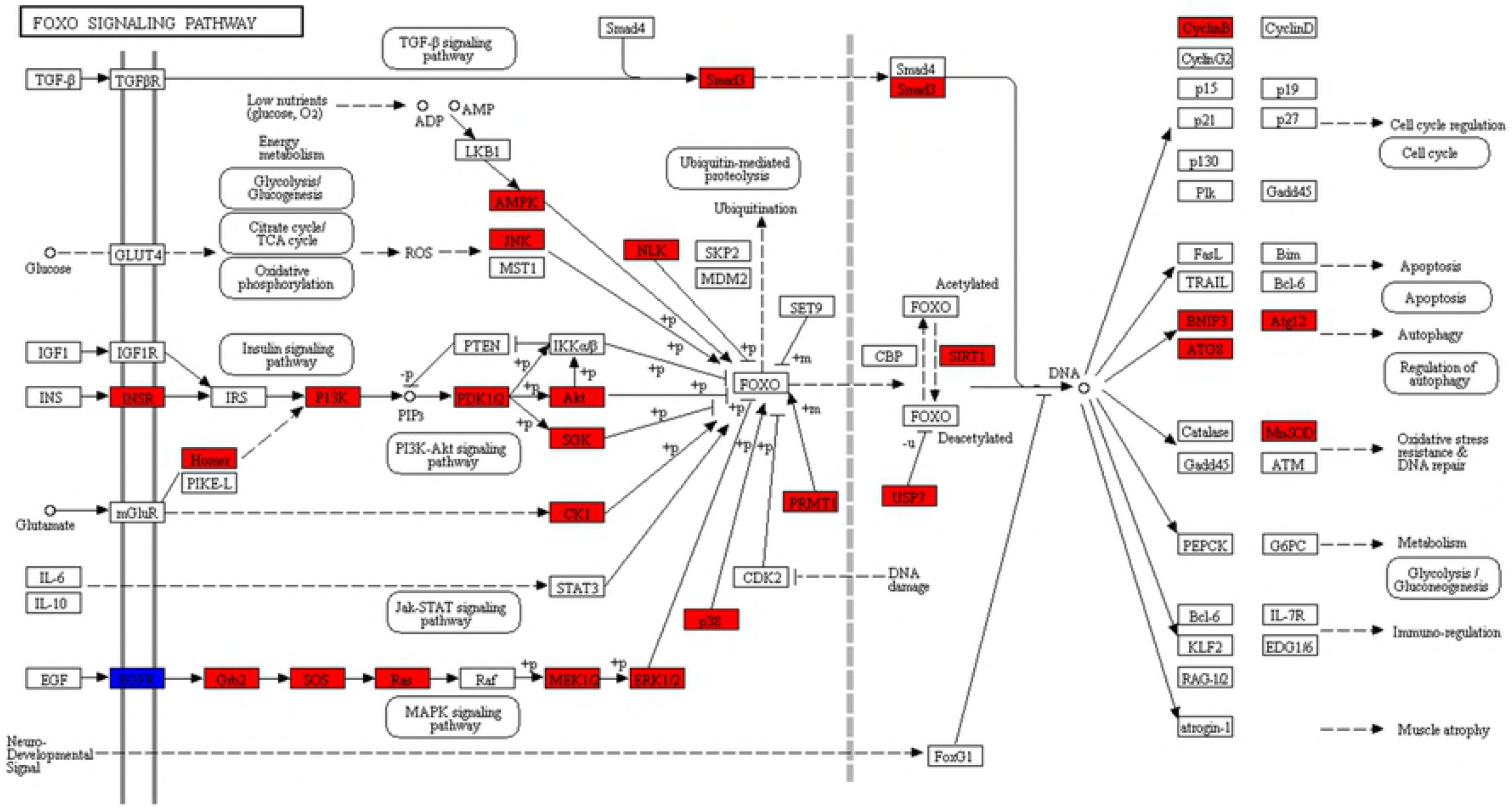
FOXO signaling pathway. Red indicates significantly up-regulated transcripts.

**Fig 7.**
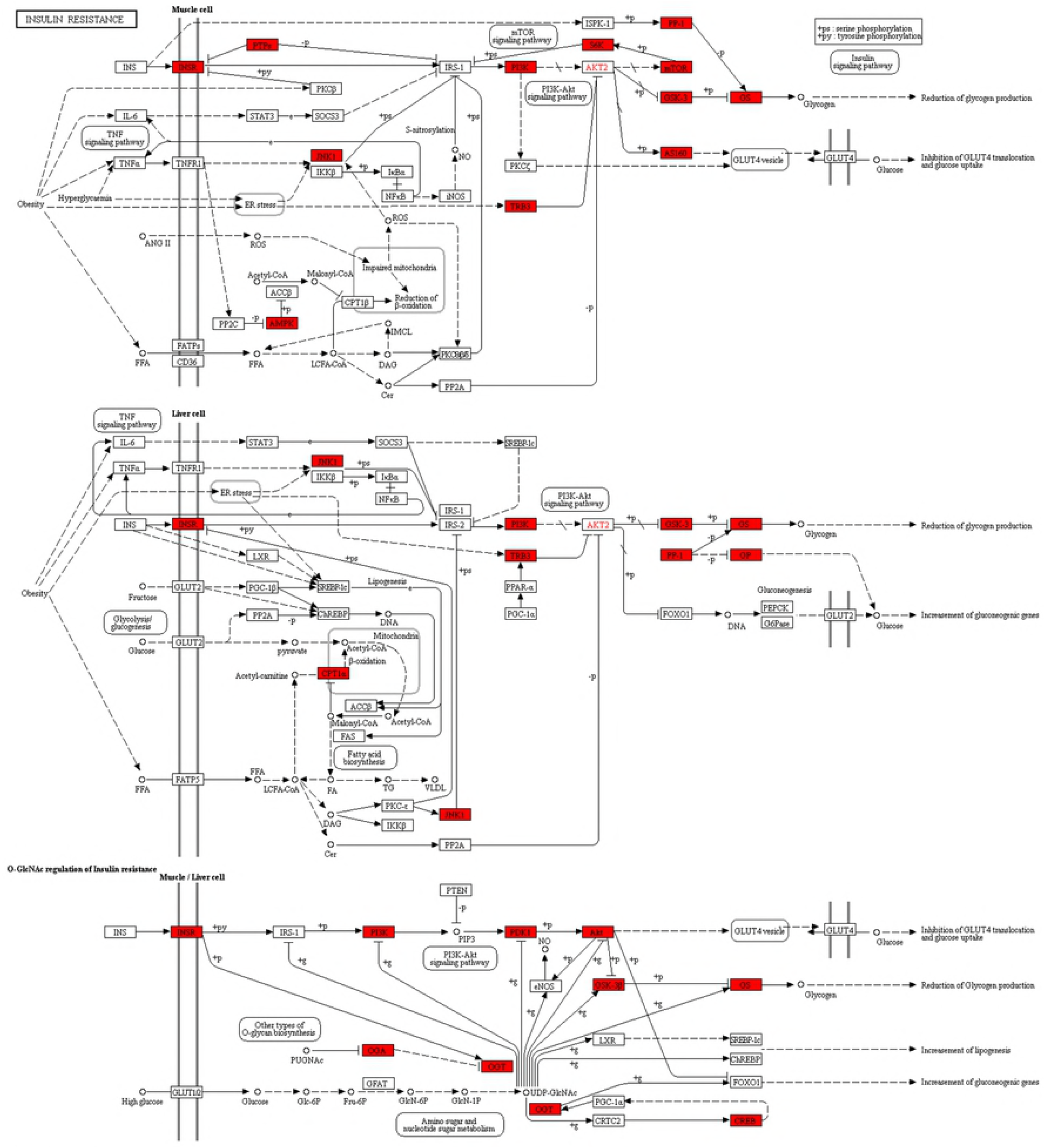
Insulin resistance pathway. Red indicates significantly up-regulated transcripts.

### Immunity- and Reproduction-Related Transcripts

The GO and KEGG analyses failed to identify any significant differences between the control and infected groups at 4 h after treatment. Therefore, in searching for genes involved in reproduction and immunity, we mainly selected significantly expressed genes in Ml6 vs CK6 group.

Of the transcripts up-regulated at 6 d after treatment, 22 were annotated as immunity-related genes (Table 4), and *defensin 3* and *PPO1* were increased the most (log_2_FC = 5.93 and Inf, respectively; ‘Inf’ indicates that the expression of the transcript in the control group was extremely low, so that the resulting ratio approached infinity). Two genes participated in Toll signaling, namely *Toll-9 receptor* and *Toll-like receptor 6,* were up-regulated in the infected group, as were *AP-1 complex subunit gamma-1, AP-2 complex subunit alpha,* and *AP-3 complex subunit mu-1, serine protease inhibitor 27A,* and *serine protease inhibitor 4-like protein.*

Seventeen reproduction-related transcripts were up-regulated, whereas both *VgA* and *VgB* were sharply down-regulated (log_2_FC = −17.82 and −18.13, respectively). Several genes related to JH metabolism, such as *juvenile hormone acid methyltransferase, juvenile hormone epoxide hydrolase-like protein 5,* and *juvenile hormone-regulated gene,* were also up-regulated in the infected group, as were *insulin metabolism related gene* and *insulin-like receptor.*

**Table 4.**
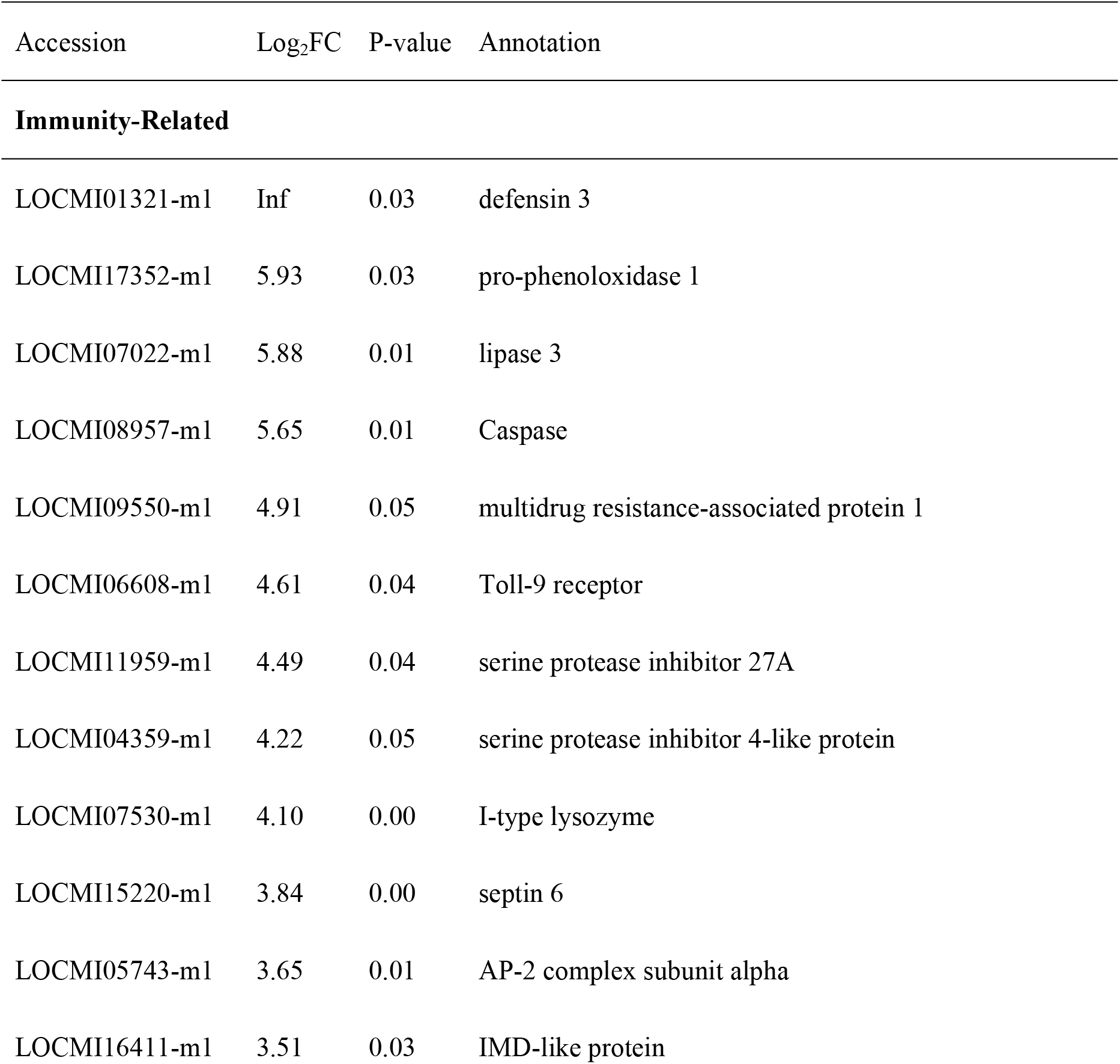
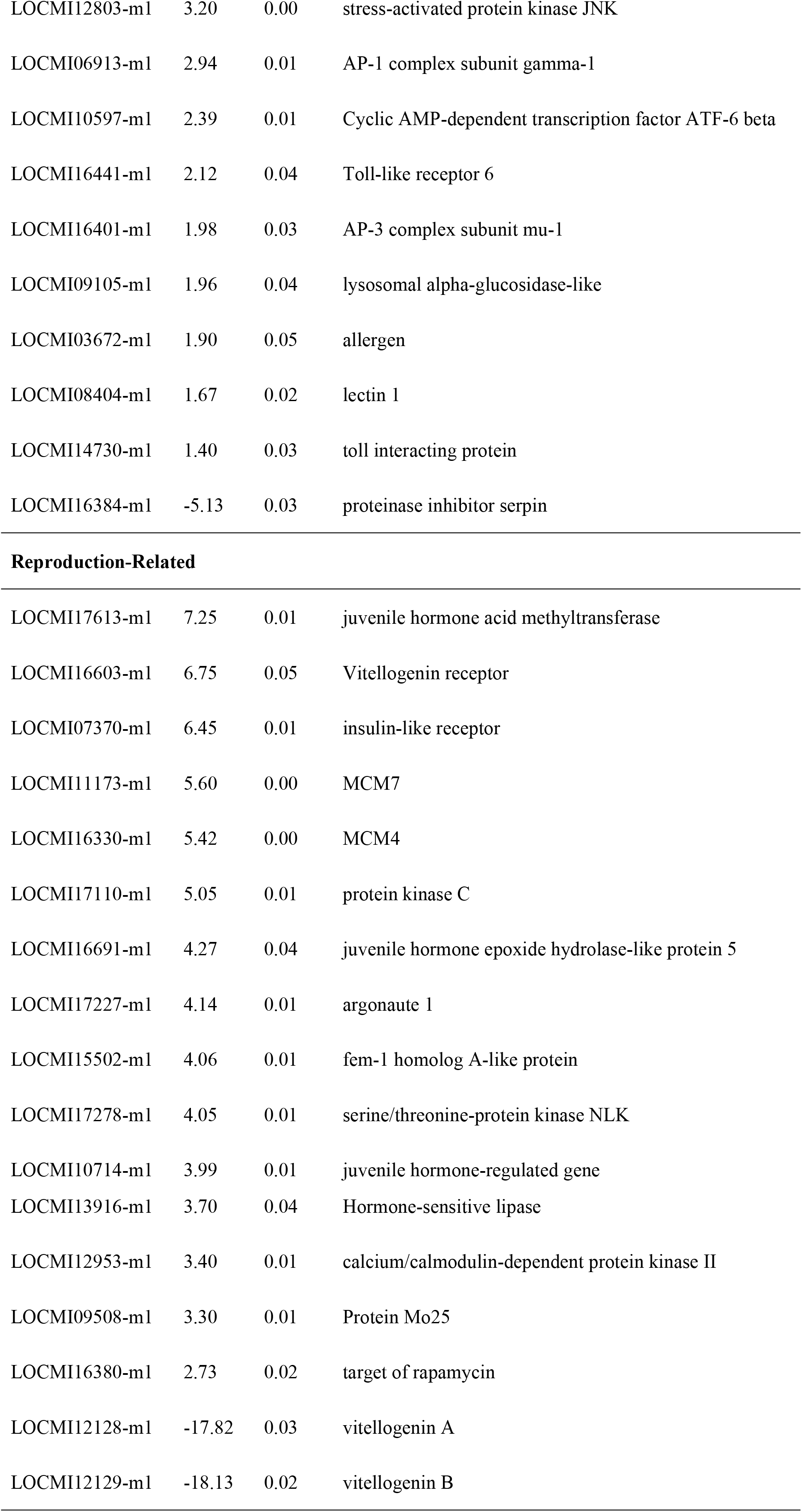
Differentially changed genes related to immunity and reproduction.

### qRT-PCR Validation of DEGs

Fifteen DEGs involved in either immunity or reproduction were randomly selected to verify the accuracy of the RNA-seq data using qRT-PCR. The qRT-PCR results validated the expression of thirteen genes (Fig 8). The differentially expression of one was not significant, while another was down-regulated. The overall trends of most of the genes were consistent, which indicated that the DGE results were reliable enough to interpret scientific problems we focused in this study (Table 5).

**Fig 8.**
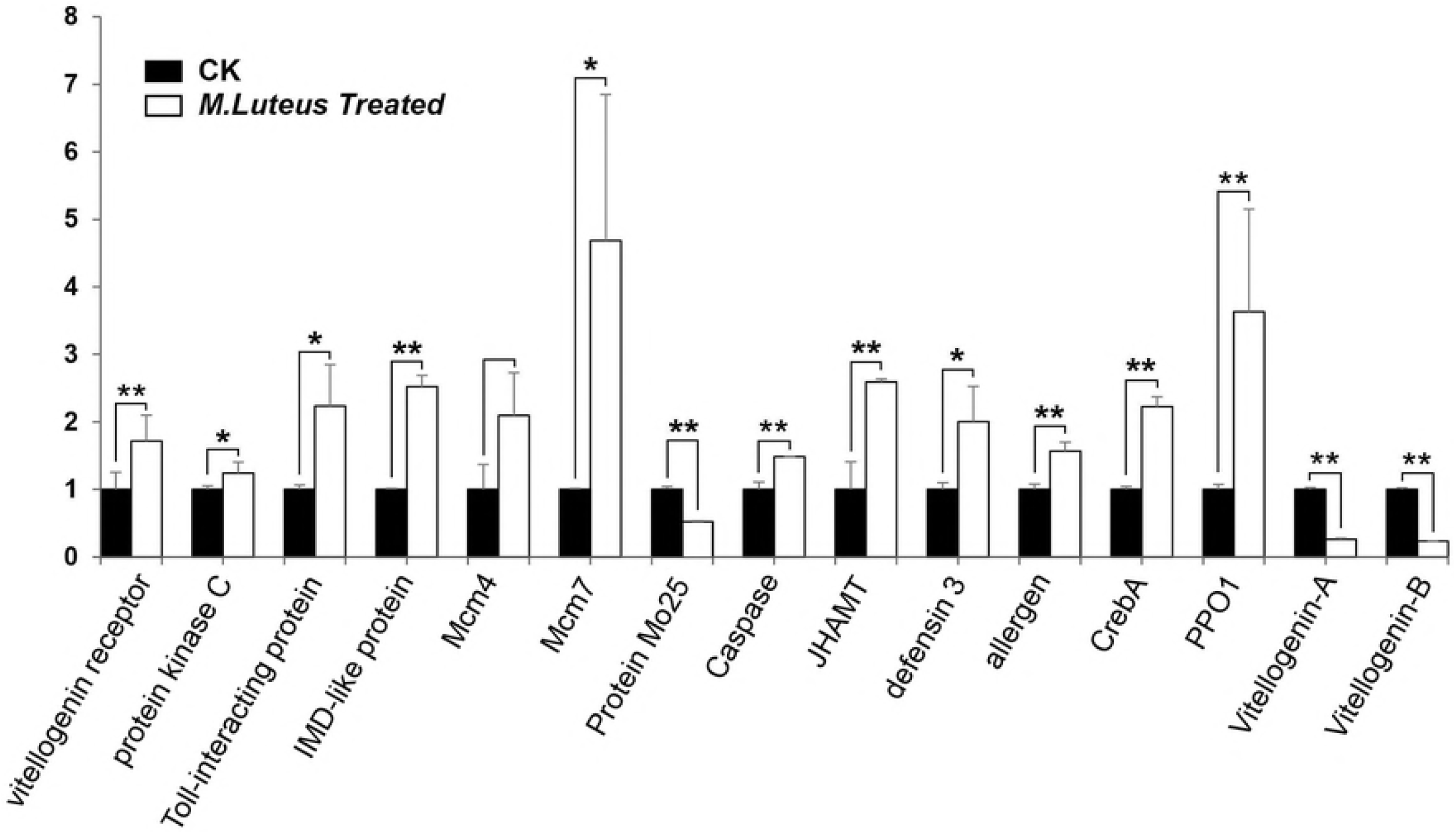
Quantitative RT-PCR analysis of 15 DEGs at 6 d after infection. “*” means P<0.05, “**” means P<0.01.

**Table 5.**
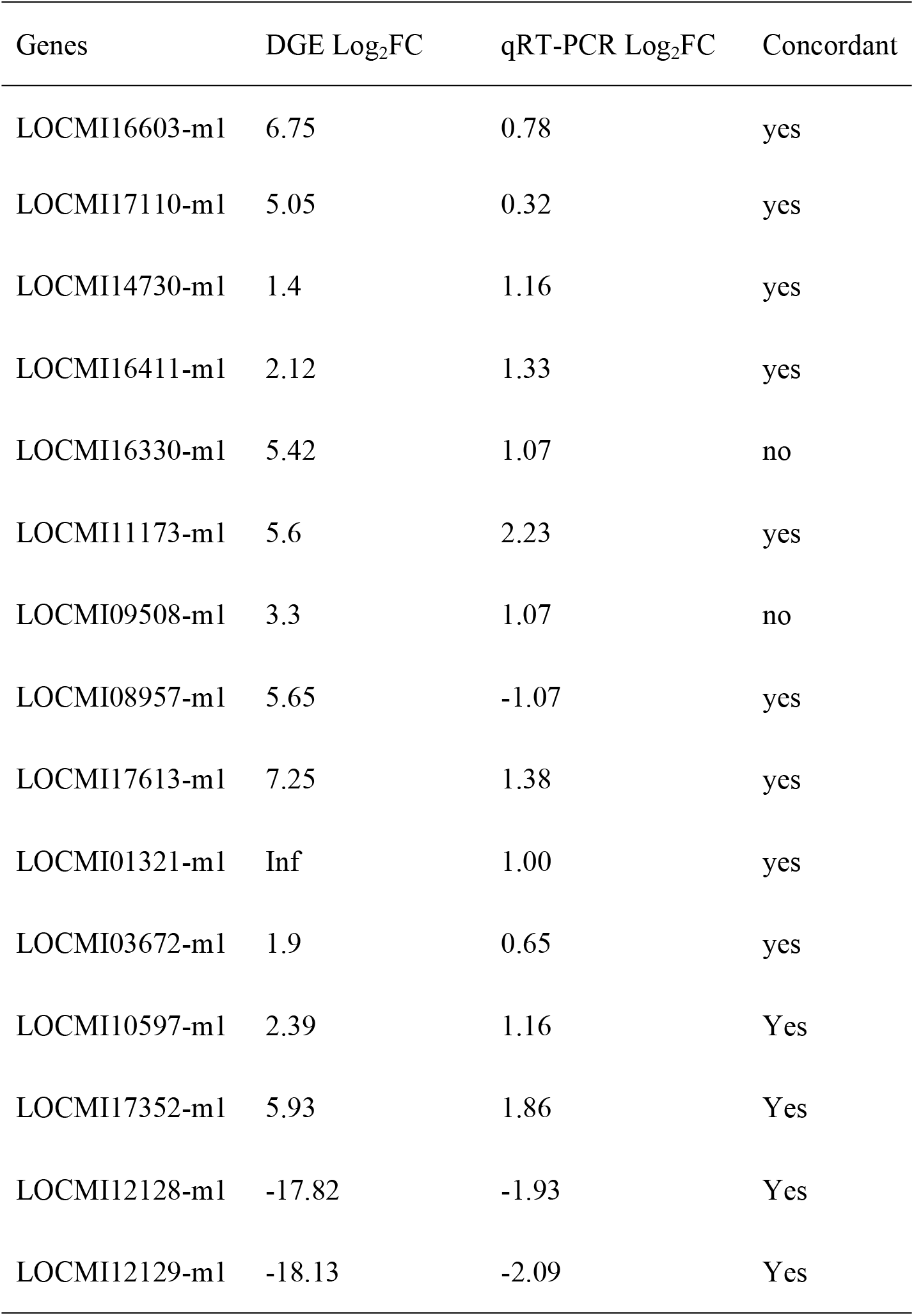
Gene models verified using qRT-PCR.

### Discussion

Resource trade-offs between immunity and reproduction may arise as a consequence of direct physiological conflicts between the two processes (Schwenkr RA et al., 2016). In the present study, the avirulent bacteria *M. luteus* was used to infect female *L. migratoria,* and the fat bodies and ovaries of the locusts were dissected at 4 h and 6 d after treatment. Female locusts develop to sexual maturity at ~5 d after eclosion, and energy and resource accumulate during this period. Significant changes in fat bodies and ovaries start at 7 to 10 d after eclosion, and varieties of ingredients in fat bodies and ovaries increased rapidly since 10 d (Xia BY et al., 1965). Bacterial infection often elicits long-term changes in global host transcription. For example, flies that have survived *M. luteus* infection remain chronically infected with ~2^10^ to 2^13^ bacteria per fly for at least 5.5 d (Troha K et al., 2018). Considering that the early response includes both an aggressive initial immune response and injury-induced transcriptional regulation (Casadevall A et al., 1999), infected insects were collected at 144 h (6 d) after treatment, with 4 h after treatment used as the control time point.

GO analysis, which was used to explore the relationship between immunity and reproduction in locusts, indicated that GO terms related catalytic activity (n = 22) and response to oxidative stress (n = 18) were the most enriched. These biological functions are the basic response of the organism to external injury stimuli. Six GO terms associated with endosome metabolism were increased significantly. In eukaryotic cells, endosomes play a central role in the regulation of fundamental processes, including nutrient uptake, immunity, signaling, adhesion, membrane turnover, and development, by regulating the reutilization or degradation of membrane components. The up-regulation of functional groups associated with endosomes indicates an increase in metabolic waste emissions and cell activity in the fat bodies and ovaries (Scott CC et al., 2014). The female locusts were at a stage where they were resisting bacterial infection and preparing for reproduction, which may have increased their metabolic activity. Lysosomes are single-membrane vesicles that release enzymes that degrade biological material (Mauvezin C et al., 2015) and, therefore, participate in waste removal. Four GO terms related to the JNK pathway, which regulates cell death in *Drosophila* and is involved in the Toll pathway (Wu C et al., 2015), were up-regulated, possibly indicating the occurrence of apoptosis in infected locust cells. GO terms for ‘innate immune response,’ ‘defense response,’ and ‘immune response’ were also up-regulated, thereby indicating that the immune response of the infected locusts was heightened. Cellular immunity related GO terms, such as ‘autophagosome,’ ‘autophagy,’ and ‘regulation of autophagy,’ were significantly up-regulated, thereby indicating that the cells were fighting against pathogenic infection (Gold KS et al., 2015). Three GO terms were involved in ovary development, including ‘ovarian follicle cell development’ (n=23), ‘ovarian follicle cell stalk formation’ (n=6), and ‘ovarian nurse cell to oocyte transport’ (n=3). The MCM complex (n=5) is a class of DNA replication genes. The JH-receptor complex has been reported to act on Mcm4 and Mcm7 to regulate DNA replication and polyploidy for vitellogenesis and oocyte maturation in migratory locusts (Guo W et al., 2014). The processes of immunity and reproduction are not separable from the nutrition metabolism of the body. Of the DEGs mapped to up-regulated GO terms, 158 and 38 were involved in immunity and reproduction, respectively, which suggested that more energy resources were allocated to bacterial defense.

The 1488 significantly altered transcripts were enriched in the KEGG pathway. The pathways related to immunity and reproduction were extracted to analyze trade-off strategies between the two processes. The top mapped pathways, namely ‘Endocytosis’ (n=83), ‘Phagosome’ (n=33), ‘Lysosome’ (n=26), and ‘Regulation of autophagy pathway’ (n=11), are involved in the elimination of infectious bacteria, viruses, and aging cells, as well as in the absorption of macromolecules in nutrient metabolism. Five DEGs were mapped to the Toll-like receptor signaling pathway, which is conserved among animals (Gay NJ et al., 2014) and can be activated by gram-positive triggers (Valanne S et al., 2011). In the present study, the *M. luteus* infection stimulated the immune response of the locusts, and the appropriate immune factors were up-regulated significantly. A total of 32 transcripts were mapped to ‘FoxO signaling pathway’. In the red flour beetle, the FoxO transcription factor can control vitellogenesis and, thereby, reproduction by negatively regulating insulin-like peptide signaling, and FoxO can also activate AMPs independently of immune pathways (Sheng Z et al., 2011). The significant up-regulation of ‘Insulin resistance’ (n=25) pathways promoted the FoxO signaling pathway (Becker T et al., 2010) and inhibited nutritional metabolism, thereby reducing or even preventing oviposition (Hsu HJ et al., 2009). This demonstrates that, at the biological pathway level, locusts allocate more energy into immune responses when faced with a trade-off between immunity and reproduction. Eighteen DEGs were mapped to the mTOR signaling pathway. It has been reported that the insulin-like peptide/TOR pathway plays a role in transducing nutritional information that regulates JH synthesis in mosquitoes and, thereby, determines the outcome of trade-offs between survival and reproduction (Perez-hedo M et al., 2013). The above analysis indicated that immune and reproductive processes are closely related to the nutritional status of individual locusts. ‘Insulin signaling pathway’ (n=4) was up-regulated by infection. Combined with the sequence information provided by the NCBI and locust genome databases (Wang XH et al., 2014), *VgA* (LOCMI12128-m1) and *VgB* (LOCMI12129-m1) were found to be down-regulated (log_2_FC = −17.82 and −18.13, respectively). Vitellogenin is the precursor of vitelline and the main egg-yolk protein in insects. It is mainly formed in fat bodies and is absorbed by oocytes through blood circulation (Chinzei Y et al., 1985). Reduced vitellogenin expression in the fat body inhibits ovarian development and oocyte maturation. In the present study, Vg expression was significantly down-regulated, which suggests that locusts, when challenged by infection, may devote the majority of resources to maintaining humoral circulation and survival and reduce investment in reproduction. Insect defensins are effector components of the innate defense system. In locusts, there are four genes that code for defensins, namely LmDEF1, 3, 4, and 5 (Lv M et al., 2016). In the present study, *defensin 3* (LmDEF3, LOCMI01321-m1) was significantly up-regulated in the infected *M. luteus* group (log_2_FC=inf). The prophenoloxidase-activating cascade is a key component of arthropod immunity. Wounding or infection triggers the release of *PPO1* through a process that involves JNK activation (Bidla G et al., 2007). In the present study, *PPO1* (LOCMI17352-m1) was up-regulated (log_2_FC = 5.93) in the infected group, which suggests that locusts are able to adapt to immune challenge. Furthermore, JH acid methyltransferase (LOCMI17613-m1) was also up-regulated (log_2_FC =7.25) in infected group. This enzyme is involved in final steps of JH biosynthesis in insects (Marchal E et al., 2011), and its up-regulation directly increases JH expression. In turn, JH plays an important role in regulating Vg synthesis and oocyte maturation (Guo W et al., 2014; Song J et al., 2014), as well as innate immunity. It has been reported in the mealworm beetle *Tenebrio molitor* that reduce of phenoloxidase, a major humoral immune effector, was mediated by JH (Rolff J et al., 2002) and that JH is a hormonal immuno-suppressor (Flatt T et al., 2008). At 6 d after emergence, locusts begin to store energy for reproduction. During this period, JH secretion increases normally but, in the infected locusts, fails to promote enough Vg secretion. The up-regulation of AMPs and *PPO1,* the markers of humoral immunity, indicates that the body has invested a lot of energy resources into infection resistance by *M. luteus.* It is inferred that locusts, in the process of balancing resource allocation between immunity and reproductive, devote more resources to eliminating infectious bacteria and maintaining their own survival than to reproduction. The molecular mechanisms of locusts trade-off of energy allocation between immunity and reproduction were represented by differentially changed GO functional groups and biological pathways. The relationship of IIS, JH, the phenoloxidase cascade, and AMP secretion in locusts should be investigated further in the future.

## Conclusion

In the present study, transcriptome analysis was used to identify *M. luteus* infection on the transcription of locust ovaries and fat bodies during the reproductive preparation period. The differentially expressed transcripts indicated that the expression of *PPO1* and *defensin 3* increased significantly, whereas the expression of *Vgs* decreased significantly, and that locusts, when faced with the trade-off of immune response and reproduction, allocate most resources to the physiological process of resistance to infection. These results provide a foundation for future studies of molecular mechanisms for immune and reproductive trade-offs in locusts and provide insight into the biological control of locusts.

## Acknowledgements

This work was supported by the National Natural Science Foundation of China (Grant Nos. 31270459 and 30970473), Project of Zhejiang Qian-Jiang Talents Program (Grant No. 2010R10093), Program for Excellent Young Teachers in Hangzhou Normal University (Grant No. JTAS 2011-01-031), and Program for Innovate Students in Hangzhou Normal University.

Sequecing was performed by Novogene (Beijing, China).

## Supplementary Material

Sup Fig 1. Lysosome pathway. Red indicates significantly up-regulated transcripts.

Sup Fig 2. mTOR signaling pathway. Red indicates significantly up-regulated transcripts.

Sup Table 1. qRT-PCR primer sequences.

Sup Table 2. Genes that were differentially expressed in infected and control locusts at 4 h after treatment.

Sup Table 3. Genes that were differentially expressed in infected and control locusts at 6 d after treatment.

Sup Table 4. Genes that were differentially expressed in control locusts at 4 h and 6 d after treatment.

Sup Table 5. Genes that were differentially expressed in infected locusts at 4 h and 6 d after treatment.

Sup Table 6. GO terms for genes that were differentially expressed in infected and control locusts at 6 d after treatment.

Sup Table 7. KEGG terms for genes that were differentially expressed in infected and control locusts at 6 d after treatment.

Sup Table 8. KEGG terms for genes that were differentially expressed in infected and control locusts at 4 h after infection.

Sup Table 9. KEGG terms for genes that were differentially expressed in the control group at 4 h and 6 d after infection.

Sup Table 10. KEGG terms for genes that were differentially expressed in the treated group at 4 h and 6 d after infection.

## Reference

1. Becker T, Loch G, Beyer M, Zinke I, Aschenbrenner AC, et al. (2010) FOXO-dependent regulation of innate immune homeostasis. Nature. 463(7279):369–373.

2. Bidla G, Dushay MS, Theopold U. (2007) Crystal cell rupture after injury in Drosophila requires the JNK pathway, small GTPases and the TNF homolog Eiger. J. Cell Sci. 120(7):1209–1215.

3. Casadevall A, Pirofski LA. (1999) Host-pathogen interactions: redefining the basic concepts of virulence and pathogenicity. Infect Immun. 67(8):3703–3713.

4. Chinzei Y, Wyatt GR. (1985) Vitellogenin titre in haemolymph of Locusta migratoria in normal adults, after ovariectomy, and in response to methoprene. J. Insect physiol. 31(6):441–445.

5. Ferrandon D, Imler JL, Hetru C, Hoffmann JA. (2007) The Drosophila systemic immune response: sensing and signalling during bacterial and fungal infections. Nat. Rev. Immunol. 7(11):862–874.

6. Flatt T, Heyland A, Rus F, Porpiglia E, Sherlock C, et al. (2008) Hormonal regulation of the humoral innate immune response in Drosophila melanogaster. J. Exp. Biol. 211(16):2712–2724.

7. Gay NJ, Symmons MF, Gangloff M, Bryant CE. (2014) Assembly and localization of Toll-like receptor signalling complexes. Nat. Rev. Immunol. 14(8):546–558.

8. Gold KS, Bruckner K. (2015) Macrophages and cellular immunity in Drosophila melanogaster. Semin Immunol. 27(6):357–368.

9. Guo W, Wu Z, Song J, Jiang F, Wang Z, et al. (2014) Juvenile hormone-receptor complex acts on mcm4 and mcm7 to promote polyploidy and vitellogenesis in the migratory locust. PLoS Genet. 10(10):e1004702.

10. Hsu HJ, Drummond-Barbosa D. (2009) Insulin levels control female germline stem cell maintenance via the niche in Drosophila. Proc. Natl. Acad. Sci. 106(4): 1117–1121.

11. Kirschman LJ, Quade AH, Zera AJ, Warne RW. (2017) Immune function trade-offs in response to parasite threats. J. Insect Physiol. 98:199–204.

12. Luo M, Li D, Wang Z, Guo W, Kang L, et al. (2017) Juvenile hormone differentially regulates two Grp78 genes encoding protein chaperones required for insect fat body cell homeostasis and vitellogenesis. J. Biol. Chem. 292(21):8823–8834.

13. Lv M, Mohamed AA, Zhang L, Zhang P, Zhang L. (2016) A family of CS alphabeta defensins and defensin-Like peptides from the migratory Locust, Locusta migratoria, and their expression dynamics during mycosis and nosemosis. PLoS One. 11(8):e0161585.

14. Marchal E, Zhang J, Badisco L, Verlinden H, Hult EF, et al (2011) Final steps in juvenile hormone biosynthesis in the desert locust, Schistocerca gregaria. Insect Biochem. Mol. Biol. 41(4):219–227.

15. Mauvezin C, Nagy P, Juhasz G, Neufeld TP. (2015) Autophagosome-lysosome fusion is independent of V-ATPase-mediated acidification. Nat. Commun. 6:7007.

16. Ma Z, Guo W, Guo X, Wang X, Kang L. (2011) Modulation of behavioral phase changes of the migratory locust by the catecholamine metabolic pathway. Proc. Natl. Acad. Sci. 108(10):3882–3887.

17. Mortazavi A, Williams BA, McCue K, Schaeffer L, Wold B. (2008) Mapping and quantifying mammalian transcriptomes by RNA-Seq. Nat. Methods. 5(7):621–628.

18. Perez-Hedo M, Rivera-Perez C, Noriega FG. (2013) The insulin/TOR signal transduction pathway is involved in the nutritional regulation of juvenile hormone synthesis in Aedes aegypti. Insect Biochem. Mol. Biol. 43(6):495–500.

19. Raikhel AS, Brown MR, Belles X. (2005) Hormonal control of reproductive processes. Comp. Mol. Insect Sci. 3: 433–491.

20. Rolff J, Siva-Jothy MT. (2002) Copulation corrupts immunity: a mechanism for a cost of mating in insects. Proc. Natl. Acad. Sci. 99(15):9916–9918.

21. Roy S, Saha TT, Zou Z, Raikhel AS. (2017) Regulatory pathways controlling female insect reproduction. Annu. Rev. Entomol. 63:489–511.

22. Scott CC, Vacca F, Gruenberg J. (2014) Endosome maturation, transport and functions. Semin. Cell Dev. Biol. 31:2–10.

23. Schwenke RA, Lazzaro BP, Wolfner MF. (2016) Reproduction-immunity trade-offs in insects. Annu. Rev. Entomol. 61:239–256.

24. Sheng Z, Xu J, Bai H, Zhu F, Palli SR. (2011) Juvenile hormone regulates vitellogenin gene expression through insulin-like peptide signaling pathway in the red flour beetle, Tribolium castaneum. J. Biol. Chem. 286(49):41924–41936.

25. Song J, Wu Z, Wang Z, Deng S, Zhou S. (2014) Kruppel-homolog 1 mediates juvenile hormone action to promote vitellogenesis and oocyte maturation in the migratory locust. Insect Biochem. Mol. Biol. 52:94–101.

26. Stearns SC. (2000) Life history evolution: successes, limitations, and prospects. Naturwissenschaften. 87(11):476–486.

27. Swevers L, Raikhel AS, Sappington TW, Shirk P, Iatrou K. (2005) Vitellogenesis and post-vitellogenic maturation of the insect ovarian follicle. Comp. Mol. Insect Sci. 1: 87–155.

28. Troha K, Im JH, Revah J, Lazzaro BP, Buchon N. (2018) Comparative transcriptomics reveals CrebA as a novel regulator of infection tolerance in D. melanogaster. PLoS Pathog. 14(2):e1006847.

29. Tufail M, Takeda M. (2008) Molecular characteristics of insect vitellogenins. J. Insect Physiol. 54(12):1447–1458.

30. Uvell H, Engstrom Y. (2007) A multilayered defense against infection: combinatorial control of insect immune genes. Trends Genet. 23(7):342–349.

31. Valanne S, Wang JH, Ramet M. (2011) The Drosophila toll signaling pathway. J. Immunol. 186(2):649–656.

32. Viljakainen L. (2015) Evolutionary genetics of insect innate immunity. Brief Funct Genomics. 14(6):407–412.

33. Wang X, Fang X, Yang P, Jiang X, Jiang F, et al. (2014) The locust genome provides insight into swarm formation and long-distance flight. Nat. Commun., 5:2957.

34. Wu C, Chen C, Dai J, Zhang F, Chen Y, et al. (2015) Toll pathway modulates TNF-induced JNK-dependent cell death in Drosophila. Open Biol. 5(7):140171.

35. Wu R, Wu Z, Wang X, Yang P, Yu D, et al. (2012) Metabolomic analysis reveals that carnitines are key regulatory metabolites in phase transition of the locusts. Proc. Natl. Acad. Sci. 109(9):3259–3263.

36. Xia BY, Guo F. (1965) The reproduction investigation of Locusta migratoria. Acta Entomologica Sinica. 14(4): 395–403.

